# cell-analysis-tools: an open-source library for single-cell analysis of multi-dimensional microscopy images

**DOI:** 10.1101/2023.03.08.531732

**Authors:** Emmanuel Contreras Guzman, Peter R. Rehani, Melissa C. Skala

**Affiliations:** Morgridge Institute for Research, 330 N Orchard St, Madison, WI 53715; University of Wisconsin-Madison, 1550 Engineering Dr, Madison, WI 53706

**Keywords:** Single-cell, image analysis, data analysis, open-source software, python

## Abstract

Single cell analysis of multi-dimensional microscopy images is repetitive, time consuming, and arduous. Numerous analysis steps are required to quantify and visualize cell heterogeneity and trends between experimental groups. The open-source community has created tools to facilitate this process. To further simplify analysis, we created a library of functions called cell-analysis-tools. This library includes functions that can streamline single-cell analysis for faster quality checking and automation. This library also includes example code with randomly generated data for dimensionality reduction [t-distributed stochastic neighbor embedding (t-SNE), principal component analysis (PCA), Uniform Manifold Approximation and Projection (UMAP)] and machine learning models [random forest, support vector machine (SVM), linear regression] that scientists can swap with their own data to visualize trends. Lastly, this library includes template scripts for feature extraction that can help identify differences between experimental groups and cell heterogeneity within a group. This library can significantly decrease user time while increasing robustness and reproducibility of results.

## 1. INTRODUCTION

Single-cell analysis of multi-dimensional microscopy images can be repetitive, time consuming, and arduous. After images have been collected, numerous analysis steps are required to quantify and visualize single-cell heterogeneity and trends between experimental groups. Some of these steps include loading images and pre-processing them, performing single-cell segmentation, loading segmentation masks, extracting features, visualizing trends, and finally inputting single-cell features into machine learning models. The open-source software community has developed a myriad of tools to perform many of these steps. In addition, we have developed a library of functions called cell-analysis-tools, to further simplify analysis for single-cell imaging. Figure 1 shows the typical steps needed for image analysis, where the highlighted sections greatly benefit from using this library. This library is tailored for single-cell analysis of multi-dimensional images, is agnostic of imaging modality, and includes functions for image processing, loading multi-dimensional images, computing mask similarity metrics, visualizing images for comparison, and functions for manipulating and visualizing fluorescence lifetime imaging (FLIM) data.

**Figure 1.**
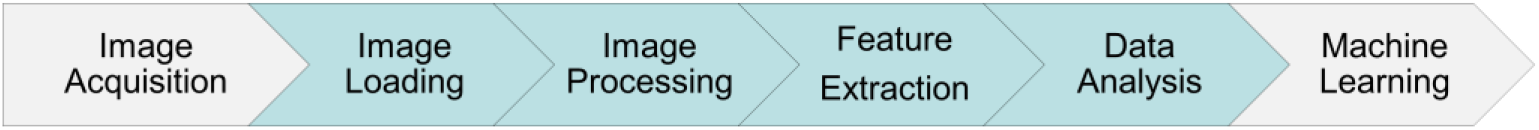
Typical steps needed for data analysis. Sections in green can greatly benefit from cell-analysis-tools.

This library was created due to the significant bottleneck of single-cell feature extraction associated with large image datasets. Each microscopy experiment is unique, so the code and functions in cell-analysis-tools are generalizable across most microscopy experiments. A central codebase enables streamlined and semi-automated analysis of new datasets and simplified quality checks for reproducible results. A template script is provided for single-cell analysis of multi-dimensional microscopy images, greatly decreasing user time while increasing the robustness and reproducibility of the results. Lastly, other templates include code examples with randomly generated data for dimensionality reduction algorithms (t-SNE, PCA, UMAP) and machine learning algorithms (random forest, SVM, linear regression) that can easily be replaced with experimental data to identify differences between experimental groups and cell heterogeneity within a group [1,2].

## 2. IMAGE ACQUISITION

Metabolic changes in cells can be visualized at a single-cell level with optical metabolic imaging (OMI). This technique uses the two-photon excitation and emission from two molecules: the reduced form of nicotinamide adenine dinucleotide (phosphate) (NAD(P)H) and flavin adenine dinucleotide (FAD), which is oxidized. NAD(P)H and FAD are already present in cells as endogenous sources of contrast [3]. The fluorescence of NADH and NADPH overlap and are collectively denoted as NAD(P)H [4]. These molecules are involved in multiple metabolic pathways including glycolysis and oxidative phosphorylation, so they can be used as a proxy for metabolic activity. Figure 2 shows a simplified diagram of these pathways including some locations where these co-enzymes are produced or consumed. To acquire images, a two-photon fluorescence microscope and a tunable laser are used to excite FAD at 890nm and collect emission with a 550/100nm bandpass filter, while NAD(P)H is excited at 750nm with emission collected through a 440/80nm bandpass filter. Time correlated single photon counting (TCSPC) electronics are used for fluorescence lifetime imaging.

**Figure 2.**
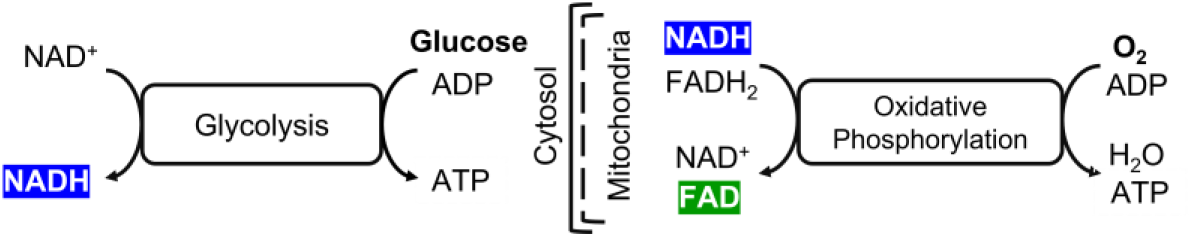
Autofluorescence imaging uses the fluorescence properties of NAD(P)H and FAD to measure cellular metabolism.

## 3. MODULES

The cell-analysis-tools library contains code to simplify intermediate steps needed to process, quantify, and analyze single-cell imaging data. The five modules that make up this library are listed in Table 1. The *Application Examples* section below highlights the use of some of these modules, functions, and template script for streamlining analysis.

**Table 1.** List of modules included in the cell-analysis-tools library.

| Module | Description |
| --- | --- |
| I/O | Loads SDT and ASC files |
| Image Processing | Pre-processes images and cleans label masks |
| Metrics | Evaluates label mask similarity or quantifies features in single-cell masks |
| Visualization | Functions to visualize images or compare generated vs. ground truth masks |
| FLIM | Functions for time-domain to frequency-domain conversions of fluorescence decays |

## 4. APPLICATION EXAMPLES

This library has been used on a variety of projects including: 1) quantifying metabolic changes in HeLa and 143B cells through time-lapse imaging during mitosis, 2) identifying differences in immune cell metabolism between healthy and graft-versus-host-disease (GVHD) for CD4^+^, CD8^+^, and CD4^+^CD8^+^ T cells, and 3) studying how *Toxoplasma gondii* alters cell metabolism. We will showcase cell-analysis-tools below using the *Toxoplasma gondii* project.

### 4.1 Toxoplasma alters cell metabolism

*Toxoplasma gondii* is the leading cause of death from foodborne illness in the United States [5]. Infection can lead to severe consequences for pregnant women and for people who are immunocompromised. In this study we aim to quantify how the metabolism of Human Foreskin Fibroblast (HFF) cells change after infection with *T. gondii.*

### 4.2 Image loading

The experiment was conducted by acquiring NAD(P)H and FAD images of HFF cells infected with mCherry-labeled *T. gondii* totaling 143 sets of images. Representative images are shown in Figure 3. HFF cells were imaged with just media or with media infected with *T. gondii.* After the images were acquired, the first step in the data analysis pipeline was to find and store paths to all the images. The data loading template script was used as a starting point for loading these images. This script recursively finds all the NAD(P)H images in a folder using regular expression, dynamically generates filenames, and finds paths to the FAD and mCherry images to form sets. These paths are then stored as a csv file for analysis.

**Figure 3.**
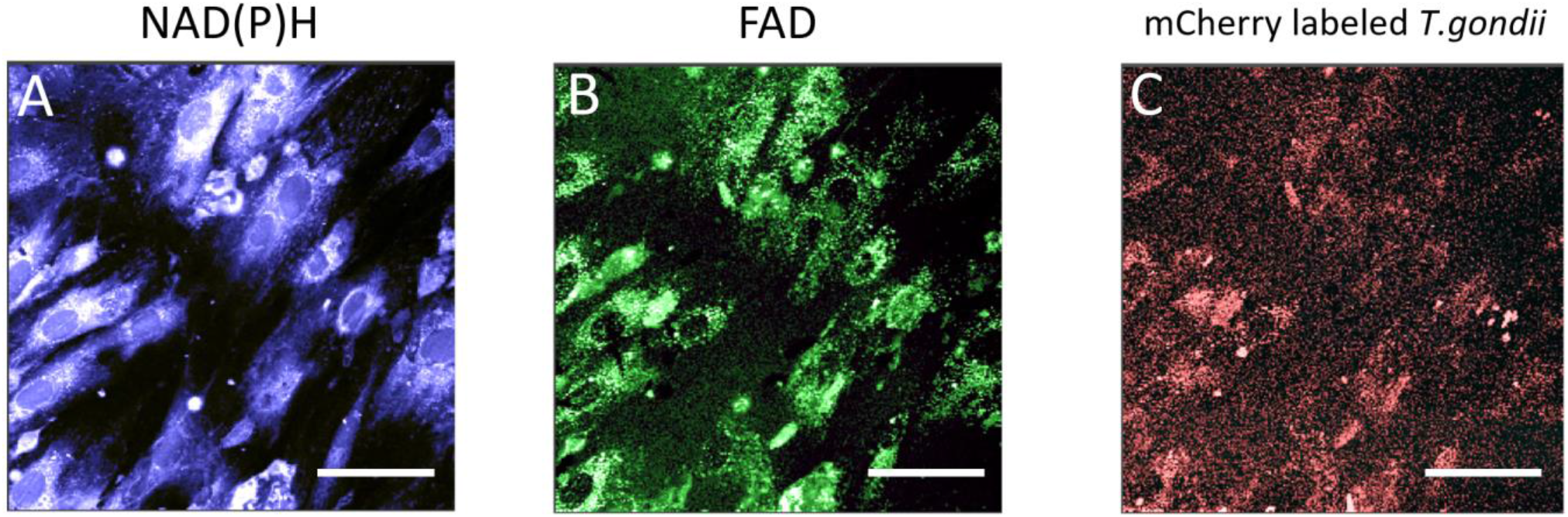
Representative set of images of A) NAD(P)H, B) FAD, and C) mCherry-labeled *T. gondii* in HFF cells. Scale-bar 40 μm.

### 4.3 Image processing

After identifying paths to all the images, these images can be pre-processed and whole-cell masks can be generated. Initial labels for masks were computed using CellProfiler [6] or Cellpose [7], then inspected and revised by a scientist using Napari image viewer [8]. *T. gondii* masks were generated using classical image processing by thresholding based on intensity as shown in Figure 4A. Masks were further cleaned up from noise and accidental duplicate labels using built-in functions that operate over each individual cell mask and remove any stray pixels that are not part of the larger connected components. Duplicate labels are removed by using the built-in functions that convert the whole-cell masks to four unique colors and then back to unique labels using the Four-Color Theorem [9]. The cleaned mask is shown in Figure 4B.

**Figure 4.**
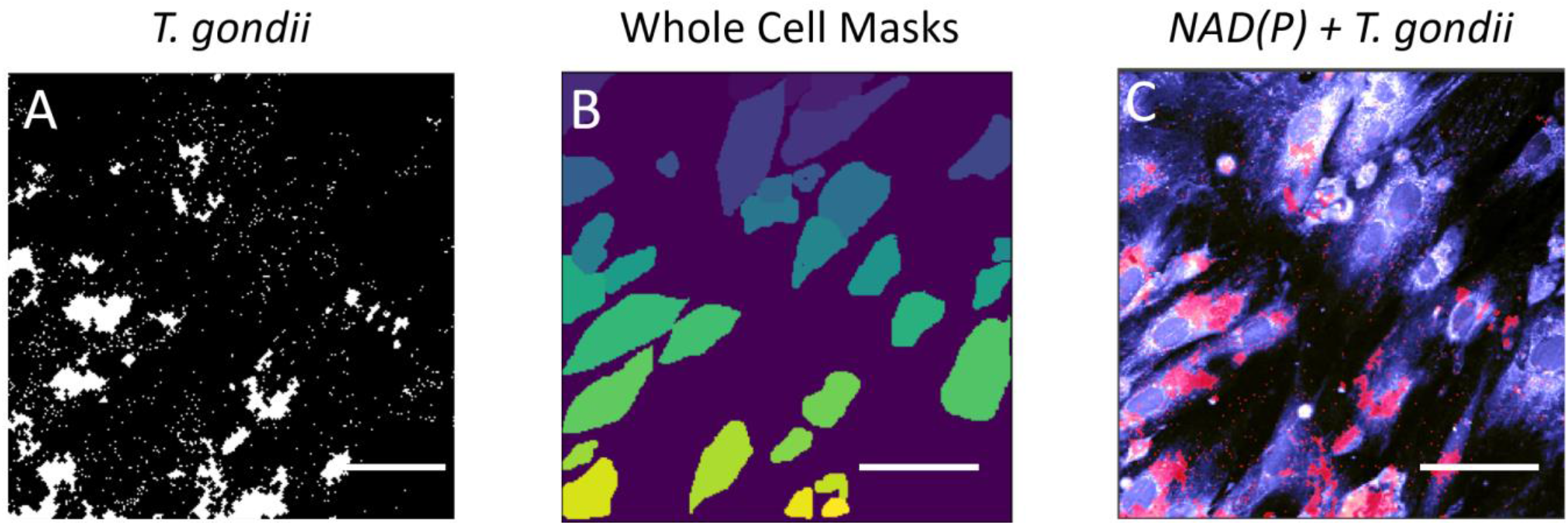
A) *T. gondii* mask, B) Whole cell mask color coded uniquely for each cell, and C) image showing overlap of NAD(P)H intensity image (blue) with *T. gondii* mask (red). Scar-bar 40 μm.

### 4.4 Feature extraction

Once the paths to all the images and masks are loaded, feature extraction is completed using a single function that takes all the processed images and masks as inputs and outputs a csv with all the features. Table 1 shows a few rows and columns of one of these csv output files. This function is an extension of scikit-image’s [10] *regionprops* function that extracts single-cell features over all images at once and computes additional features, like standard deviation or redox ratio [11], that the original function does not provide.

**Table 1.** The first three rows of an example CSV, showing a subset of features quantified for each cell.

| Filename | mask<br>label | NAD(P)H Intensity<br>mean | NAD(P)H Intensity<br>stdev | NAD(P)H a1<br>mean | NAD(P)H a1<br>stdev | NAD(P)H t1<br>mean | NAD(P)H t1<br>stdev |
| --- | --- | --- | --- | --- | --- | --- | --- |
| 20190409_control_Tgondii_001 | 1 | 468.18 | 137.82 | 74.00 | 4.41 | 492.42 | 76.45 |
| 20190409_control_Tgondii_001 | 2 | 242.90 | 106.74 | 73.99 | 7.77 | 506.19 | 139.08 |
| 20190409_control_Tgondii_001 | 3 | 396.23 | 169.73 | 72.53 | 5.73 | 481.51 | 92.55 |

### 4.5 Data analysis

With the extracted features, trends in the data can be analyzed and visualized. For example, the template code can plot a PCA and UMAP of the data to visualize any clustering by condition (Figure 6) [1,2]. Here, the uninfected and infected cells show clear separation.

**Figure 6.**
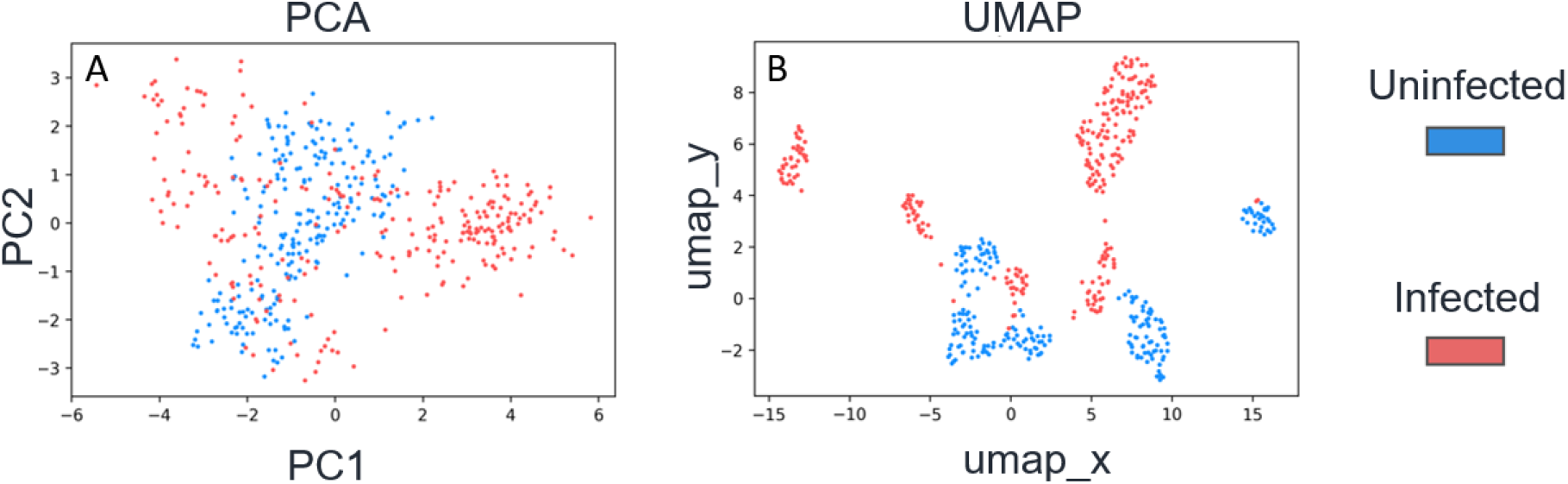
(A) PCA and (B) UMAP generated using the template script. Clustering is observed between the uninfected and the infected conditions in both the PCA and UMAP.

## 5. CONCLUSION

There are many steps required to analyze single-cell imaging data. The cell-analysis-tools library has many advantages including: 1) automated feature extraction from images for single-cell analysis, 2) fast quality control checks of data, 3) increased reproducibility and robustness of results, 4) generalized code base can be used broadly across projects, and 5) bug fixes can be easily propagated across projects that use the same codebase.

## CODE

The code carries a GPL-3 license and is publicly available on GitHub at https://github.com/skalalab/cell-analysis-tools along with the documentation https://cell-analysis-tools.readthedocs.io.

## ACKNOWLEDGEMENTS

Special thanks to all the Skala lab members: Rupsa Datta, Kayvan Samimi, Alexa Heaton, Danielle Desa, Amani Gillette, Dan Pham, Anne-Sophie Mancha, Kasia Wiech, Jeremiah Riendeau, Heather Esser, Andrea Schiefelbein, Skala Lab Undergraduates and High Schoolers, Former lab members as well as Gina Gallego-Lopez and Laura Knoll from the Knoll lab.

